# Robust Hierarchical Density Estimation and Regression for Re-stained Histological Whole Slide Image Co-registration

**DOI:** 10.1101/565564

**Authors:** Jun Jiang, Nicholas B. Larson, Naresh Prodduturi, Thomas J. Flotte, Steven N. Hart

## Abstract

For many disease conditions, tissue samples are colored with multiple dyes and stains to add contrast and location information for specific proteins to accurately identify and diagnose disease. This presents a computational challenge for digital pathology, as whole-slide images (WSIs) need to be properly overlaid (i.e. registered) to identify co-localized features. Traditional image registration methods sometimes fail due to the high variation of cell density and insufficient texture information in WSIs – particularly at high magnifications. In this paper, we proposed a robust image registration strategy to align re-stained WSIs precisely and efficiently. This method is applied to 30 pairs of immunohistochemical (IHC) stains and their hematoxylin and eosin (H&E) counterparts. Our approach advances the existing methods in three key ways. First, we introduce refinements to existing image registration methods. Second, we present an effective weighting strategy using kernel density estimation to mitigate registration errors. Third, we account for the linear relationship across WSI levels to improve accuracy. Our experiments show significant decreases in registration errors when on matching IHC and H&E pairs, enabling subcellular-level analysis on stained and re-stained histological images. We also provide a tool to allow users to develop their own registration benchmarking experiments.

## Introduction

Histological assessment is a critically important feature in the diagnosis, prognosis, and treatment of disease. Tissues are frequently viewed under a traditional light microscope after some type of tissue staining – most commonly hematoxylin & eosin (H&E) and immunohistochemistry (IHC). High-throughput whole-slide scanners are now enabling quantitative assessments of histologic features from whole-slide images (WSIs).

While exciting, these new advances in digital pathology also bring new challenges. One such challenge is co-locate features from the same tissue after it has been stained, imaged, washed, stained again, and imaged again. This type of procedure is common when features need to be compared at the sub-cellular level. Unlike serial sectioning, where pieces of tissue are cut sequentially, re-staining does not suffer from the potential distortions associated with architectural changes from differences in the tissue sections. Serial sections are commonly used to identify characteristics of a tumor (e.g. estrogen receptor positivity in breast cancer) but not for the same cell.

From a computational perspective, this means the re-staining scenario is more suitable for rigid image registration, while serial sectioning is more suitable for non-rigid image registration. Most of the previous work in the WSI registration field has been focused on aligning serial sections of tissues for cross-sectional observations (e.g., 3-dimensional tissue visualization) [1-5]. Methods such as those proposed by Mueller et al. [6] use a deformable multi-modal WSI registration technique are well suited for serial section registration. However, using deformable registration does not explicitly model the fact that pixels in multi-modal images do not belong to the same cell, which may introduce errors into downstream analysis. Here, we only focus on the re-stained histological WSI co-registration use case.

The need for a rigid image registration is based on the fact that re-stained tissues have a fixed relative position to the slide glass, meaning there should be some (x, y) shifting, but minimal rotation. Several state-of-the-art rigid image registration methods could potentially be used to handle this problem. Scale Invariant Feature Transform (SIFT) [16] is one of the most popular “key points”-based image registration method, in which strongly differentiable pixel areas are detected, filtered, and matched based on their local features. Key points can be filtered based on the differences in slope between matched key point pairs. In other words, lines connecting matched pairs of key points in the stationary and corresponding floating image should be parallel. Enhanced Correlation Coefficient (ECC) [14] has also been shown to successfully align single channel images to generate multi-channel images. Finally, Fast Fourier Transform (FFT) [15] image registration methods are generally insensitive to translation, rotation, scaling, and noise, with the added benefit of being computationally efficient. Each of these methods can return a success rate and similarity score (as a measure of alignment confidence) for each registration by calculating the image similarity after image registration.

The computational cost of directly aligning gigapixel WSIs can be prohibitively high, so a smaller “image patch”-based method is often required. By virtue of their smaller size, patch-based methods do not contain as much texture information to use for registration, which can ultimately lead to poor image alignments. To address these issues, Rosetti et al.[4] proposed a method to align images at lower resolution level and propagate the registration offset to high resolutions. However, they assume that tissue contours are relatively unchanged in the grayscale space – which is not the case between H&E and IHC. Obando’s method [7] refines the common information between downsampled images before attempting registration, whereas Trahearn’s method [2] filters regions based on entropy, thresholds, and texture features.

There are several drawbacks to all these methods. First, WSIs with different stains may look quite different not only in hue and contrast, but also different in local details, because H&E stains both nucleus and cytoplasm, while IHC only highlights the location of the target protein. This can cause methods that rely on finding key points (key point matching) to fail, yielding more failures to align as too few suitable correlations in the images can be found. The other major problem is that the registration errors in low-resolution images will be magnified dozens of times if the transformation is directly applied to high-resolution levels. Therefore, a more accurate way to leverage the information from lower resolution to high resolution is needed.

In this paper, we proposed a robust strategy for WSI co-registration which can precisely and efficiently align re-stained WSIs of the same tissue sections. Our approach leverages image patches from the information-rich low resolution layers, weights their offsets by kernel density estimation (KDE), and then regresses against the hierarchical nature of WSIs to refine the offset parameters between two WSIs. Our results suggest that simple rigid image registration that incorporates the unique structure of WSI file hierarchy is well suited to address the re-stained WSI registration tasks.

## Method

The main workflow of our registration strategy has been illustrated in **Figure 1**. There are multiple scaled image levels in WSIs. We extracted image patches from the top three image levels (the lowest resolution), and adopted Fourier transforms-based image registration to align H&E and IHC pairs to each level. Since there are linear relationships between image levels, we proposed a weighted hierarchical linear regression method to get the optimal registration offset and rotation angle. Generally, our approach consists of three main parts: 1) image patch-based registration; 2) KDE; 3) hierarchical resolution regression

**Figure 1.**
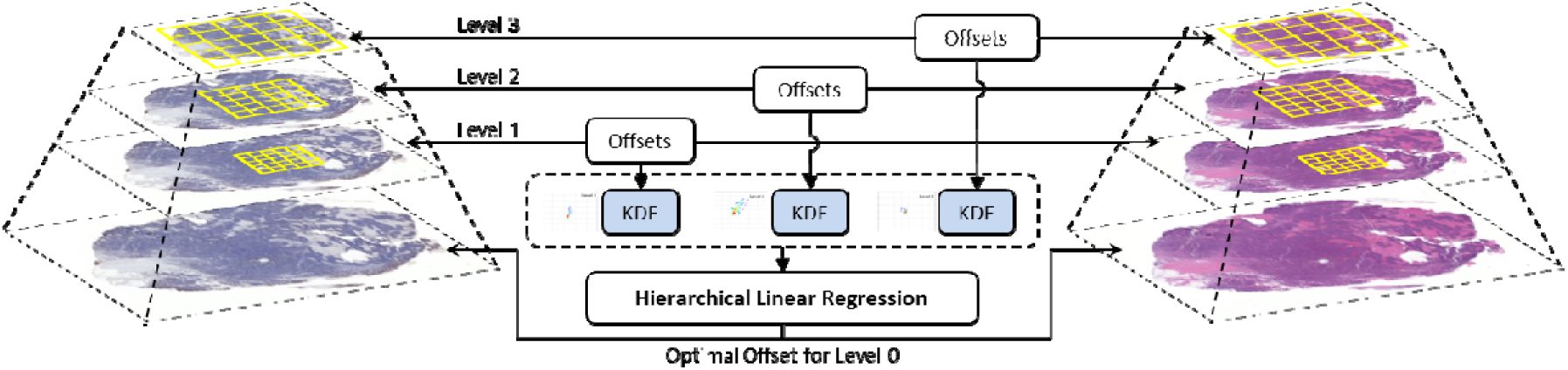
An overview of our proposed method. The first step is get raw registration result from top three whole slide image levels. The second step is adopting Kernel Density Estimation to weight the raw registration. The last step is using hierarchical linear regression to get the optimal co-registration for whole slide images.

### 1. Image patch-based registration

In order to leverage low-level image features in WSIs, we introduced Fourier transform based image registration [8]. The goal of image registration is to determine a transformation that maximizes the similarity between two images. The image registration problem can be formulated as: I′ = T · I, in which I denotes the original image, and I′, denotes the transformed image. In our histological image registration task, there is no scale and affine transformation, because we want to align the re-stained slides under the same magnification. For a two-dimensional image registration problem, the transformation matrix T can be written like below.

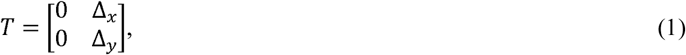

where Δ_*x*_ and Δ_*y*_ denote *x* and *y* offsets, respectively. If *f*_2_(*x,y*) is a translated and rotated replica of *f*_1_(*x,y*) with translation (Δ_*x*_, Δ_*y*_), then

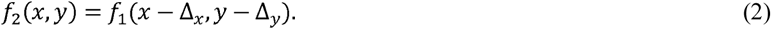

According to the Fourier translation and rotation properties, transforms of *f*_1_ and *f*_2_ are related by

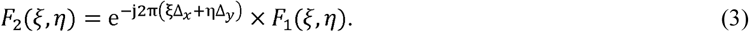

The cross-power spectrum of two images is defined as

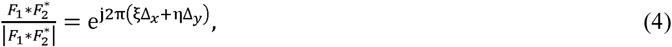

where 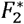 is the complex conjugate of *F*_2_. The shift theorem guarantees that the phase of the cross-power spectrum is equivalent to the phase difference between the images. By taking the inverse Fourier transform of the representation in the frequency domain, we have an impulse function that is approximately zero everywhere except at the displacement that is needed to optimally register the two images. We choose the “imreg_dft” library [9] as the basis of our framework, which is an implementation of discrete Fourier transformation based image registration [10].

### 2. Kernel Density Estimation

Fourier transforms-based image registration method provides a straightforward approach to align two image patches. We initially tried to directly register pathological image pairs together in highest resolution with this method, and found that most pairs of patches can be aligned well, but several failed due to limited texture information for use in defining key points. To overcome this limitation, offsets were computed from multiple patches sampled from the same WSI and at the same resolution. At first, we simply computed the mean offset in the x and y planes, but found this approach was highly variable (**Figure 2**). Instead, we introduced a KDE algorithm [11] to estimate the most possible offset for a robust image alignment.

**Figure 2.**
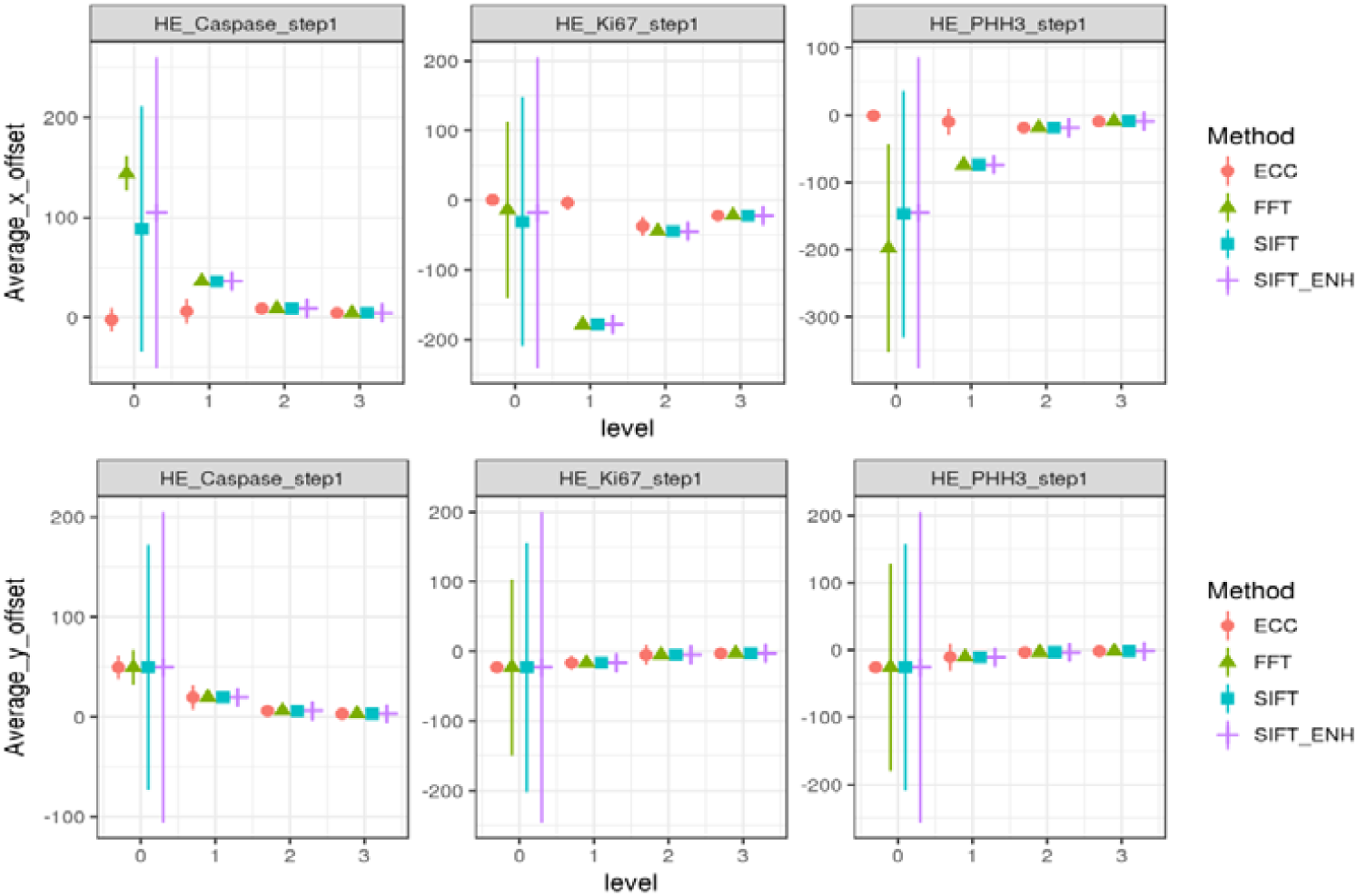
Registration offsets (mean and 1 standard deviation) including four different methods on four image levels. The first row represents the horizontal offset; the second row represents the vertical offset. Each column represents the offset between an H&E-IHC WSI pair. Ideally, the error should be clustered toward zero. The high variance in level 0 indicates frequent registration failures.

KDE is a non-parametric method to estimate the probability density function of a random variable, which can be defined as follows:

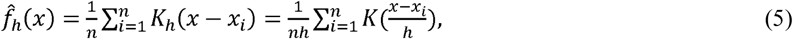

where *x*_*i*_ is a sampled observation, *n* is the sample size, 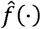 is an estimate the probability density function of *x, K*(·) is a non-negative kernel function, and h > 0 is the bandwidth smoothing parameter. This formula can also estimate image rotation, *θ*. In our study, the registration offset (*x, y*) is a two-dimensional variable, so the KDE with a Gaussian kernel can be defined as:

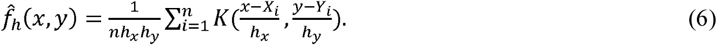

We apply the Scott’s rule [12] to define bandwidth, and let the bandwidth be the same on both x and y directions, such that

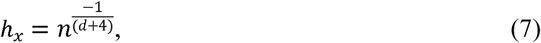

where *d* is the dimensionality of each data point. In our study, we let *h*_*x*_ = *h*_*y*_ = *h*_*xy*_ = *h*_*yx*_ because we have no prior knowledge about the offset distribution.

Since there is no bias in the choice of image patches during image registration, we can arbitrarily use Gaussian approximation as our kernel function:

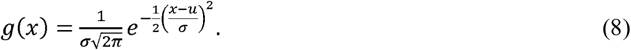

With a Gaussian kernel, the estimator is the weighted sum of the bivariate normal densities:

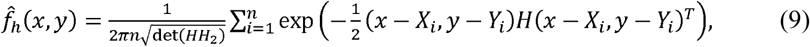

where (*X, Y*) denotes the registration offsets. The matrix *H* is the covariance matrix expressed in terms of bandwidth, as defined in formula (7).

We can regard the density estimates as the measures of confidence to weight the offset and angle for image pairs. Density estimation for every registration offset can be denoted as 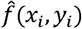, and normalized the confidence to [0,1]. By applying KDE, we mitigate the influence of registration errors by down-weighting their contribution to estimates of overall registration offsets and rotation angles.

### 3. Hierarchical Resolution Regression

There are several levels in a WSI (**Figure 1**), which are defined by the multi-resolution pyramidal data structure. The histological image pairs could be very different in details at lower levels (with high resolution), but appear similar in higher levels (with low resolution). This phenomenon is especially common in consecutive tissue sections. If the registration is performed only at the lowest resolution, the image registration may fail due to the weak texture information in some areas. Considering there are many image patches available at each image level, we can develop a robust method by calibrating the registration offsets across different image levels.

To find the pixel correlations across image levels, we investigated the relationship of registration offsets for two image levels. If we denote a pixel offset p(x,y) in level 1 with image resolution R_1_, and denote the corresponding pixel offset p(x′,y′) in level 2 with image resolution R_2_, and the downsampling factor between image levels is known and fixed, it follows that pixel correlations can be defined as:

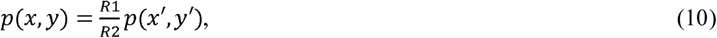

So, if we apply image registration on two different image levels, in ideal conditions, the linear regression of offsets should be a line that goes through the origin. A weighted linear regression strategy can be used to adaptively learn the exact linear relationship. Therefore, the cost function of the regression problem can be interpreted as:

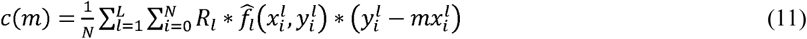

where R_*l*_ ∈ {0.25,0.25,0.5} denotes the resolution ratio of current image level to lower image level, *L* ∈ [1,3] denotes the image levels, *N* represents the number of image patches used at each level, and 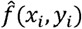 is calculated from formula (9). By minimizing this cost function, the optimal slope *m* for cross-level image registration can be determined.

## Experimental Data

### A. Data Description

We collected 30 WSI pairs (90 slides in total) of an H&E and an IHC image of the same tissue section (H&E vs Caspase3, KI67 and PHH3). OpenSlide [13], a public C library was introduced to extract image patches from a specific coordinate and image level. There are four image levels in each WSI: levels 0-3, where level 0 denotes the original image with highest resolution and level 3 denotes the down-sampled image with lowest resolution. The resolution ratio of image levels for our dataset is 1:4:16:32 (from level 3 to level 0), with a pixel size is 0.25µm at the highest resolution.

For destaining H&E sections, the slides are placed in xylene until the coverslips float off the slide. The slides are placed in three containers of xylene for 5 minutes each, followed by two containers of absolute alcohol for 1 minute each, three containers of 95% alcohol for one minute each and one container of 70% alcohol for one minute. The slides are rinsed in tap water until clear.

IHC staining was performed at the Pathology Research Core (Mayo Clinic, Rochester, MN) using the Leica Bond RX stainer (Leica). FFPE tissues were sectioned at 5 microns and IHC staining was performed on-line. Slides were retrieved for 20 minutes using Epitope Retrieval 2 (EDTA; Leica) and incubated in protein block (Rodent Block M, Biocare) for 30 minutes. The phospho-Histone H3 primary antibody (Catalog #9701; Cell Signaling) was diluted to 1:50 in Bond Antibody Diluent (Leica). The cleaved-Caspase 3 primary antibody (Rabbit Polyclonal – Catalog 9661L; Cell Signaling) was diluted to 1:200 in Bond Antibody Diluent (Leica). The Ki-67 primary antibody (Clone MIB-1, Dako) was diluted to 1:300 in Bond Antibody Diluent (Leica). All primary antibodies were incubated for 15 minutes.

The detection system used was Polymer Refine Detection System (Leica). This system includes the hydrogen peroxidase block, post primary and polymer reagent, DAB, and Hematoxylin. Immunostaining visualization was achieved by incubating slides 10 minutes in DAB and DAB buffer (1:19 mixture) from the Bond Polymer Refine Detection System. To this point, slides were rinsed between steps with 1X Bond Wash Buffer (Leica). Slides were counterstained for five minutes using Schmidt hematoxylin and molecular biology grade water (1:1 mixture), followed by several rinses in 1X Bond wash buffer and distilled water, this is not the hematoxylin provided with the Refine kit. Once the immunochemistry process was completed, slides were removed from the stainer and rinsed in tap water for five minutes. Slides were dehydrated in increasing concentrations of ethyl alcohol and cleared in 3 changes of xylene prior to permanent coverslipping in xylene-based medium.

All breast cancer specimens were excess tissue at surgical resection and were obtained with IRB approval. The specimens were fixed in buffered formalin following the ASCO/CAP guidelines for HER2 testing for between 6 to 72 hours.

### B. Ground Truth Annotation

There is no general quantified evaluation metric for image registration of multiple WSIs. To evaluate the registration accuracy, we developed an image visualization tool to manually align two WSIs. Ground truth was obtained by adjusting a floating image’s offset with respect to a fixed image (**Figure 3**). The ground truth of image registration for our dataset was determined by comparing the details at the same position after careful manual adjustment of the floating image. We have made this tool freely available for others to use in their own projects. ^1^https://github.com/smujiang/Re-stained_WSIs_Registration

**Figure 3.**
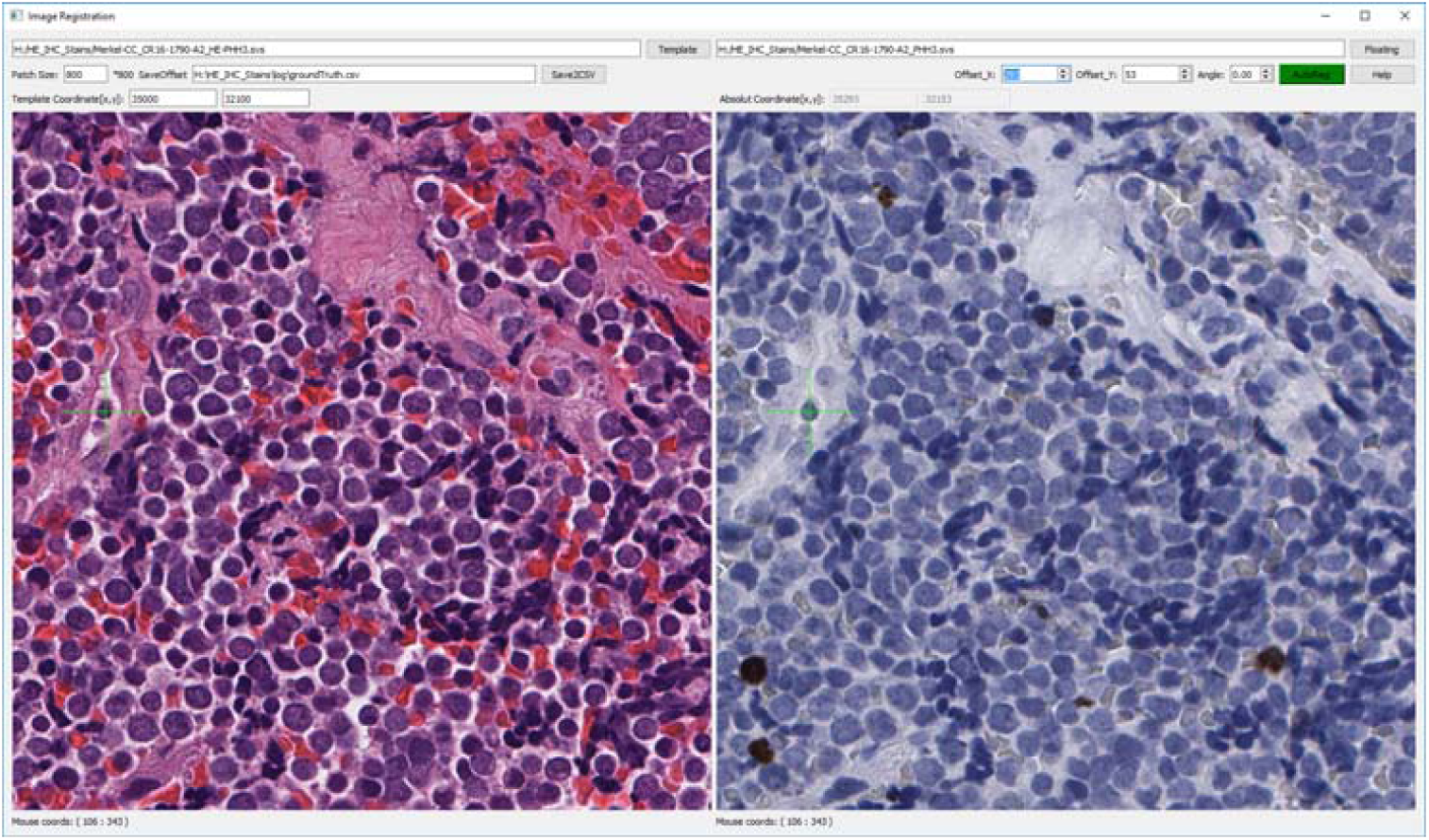
Screen shot of WSI registration tool. After loading WSIs, the image on the right can be shifted and rotated by adjusting the x and y offset positions. A green cross is attached to mouse cursor, so that details of two images at the same location can be compared easily. Source code of this tool can be found at our GitHub^1^.

## Experiments and Results

Our experiments were conducted as illustrated in the workflow (**Figure 1**), which consists of three main steps: 1) raw registration, 2) KDE weighting, and 3) hierarchical linear regression.

### 1. Raw Registration

For each pair of H&E-IHC WSIs, an initial offset was obtained by align the thumbnails of the WSIs. Image brightness thresholding was used to locate content rich area, from which we randomly sample image patches for raw registration. We co-registered pairs of H&E-IHC patches on all WSI levels using three rigid registration methods: ECC [14], FFT[15], and SIFT [16]. To improve registration accuracy, we also tried to add an extra filter to distill the matched key points detected by RANSAC [17] in traditional SIFT output, which is denoted as SIFT-ENH. The extra filter keeps key points that share a similar slope. All implementations of these methods are based on the OpenCV API [18]. We applied FFT together with these three methods to our dataset, the statistical features of registration results are shown in **Figure 2**. ECC, FFT, SIFT and SIFT_ENH all performs well on lower resolutions (levels 2-3), but poor at level 0.

### 2. KDE Weighting

To reduce the variance in the predicted offsets, we attempted two different types of procedures: weighting based on raw registration score and kernel density estimation (KDE). All the image registration methods assessed here return a score to measure the success of registration. However, we found that these scores do not always reflect the registration success for many patch-level registrations. The reason might come from the fact that H&E and IHC images are very different in hue and texture, and some have sparse unique local details. To reveal this phenomenon, we show an example (**Figure 4**, top row) by drawing the registration offsets in two dimensional coordinates, where each registration is represented as a dot, colored by confidence score. High scores mean the registration algorithm has high confidence in its offset estimation. We found that offsets close to the ground truth are not necessarily those with higher scores. This deficiency implies that confidence score based on image similarity cannot effectively measure the registration accuracy, and a more efficient way should be adopted. Hence, we show the KDE algorithm is more accurate at predicting the higher scoring alignments than native scoring methods (**Figure 4**, bottom row).

**Figure 4.**
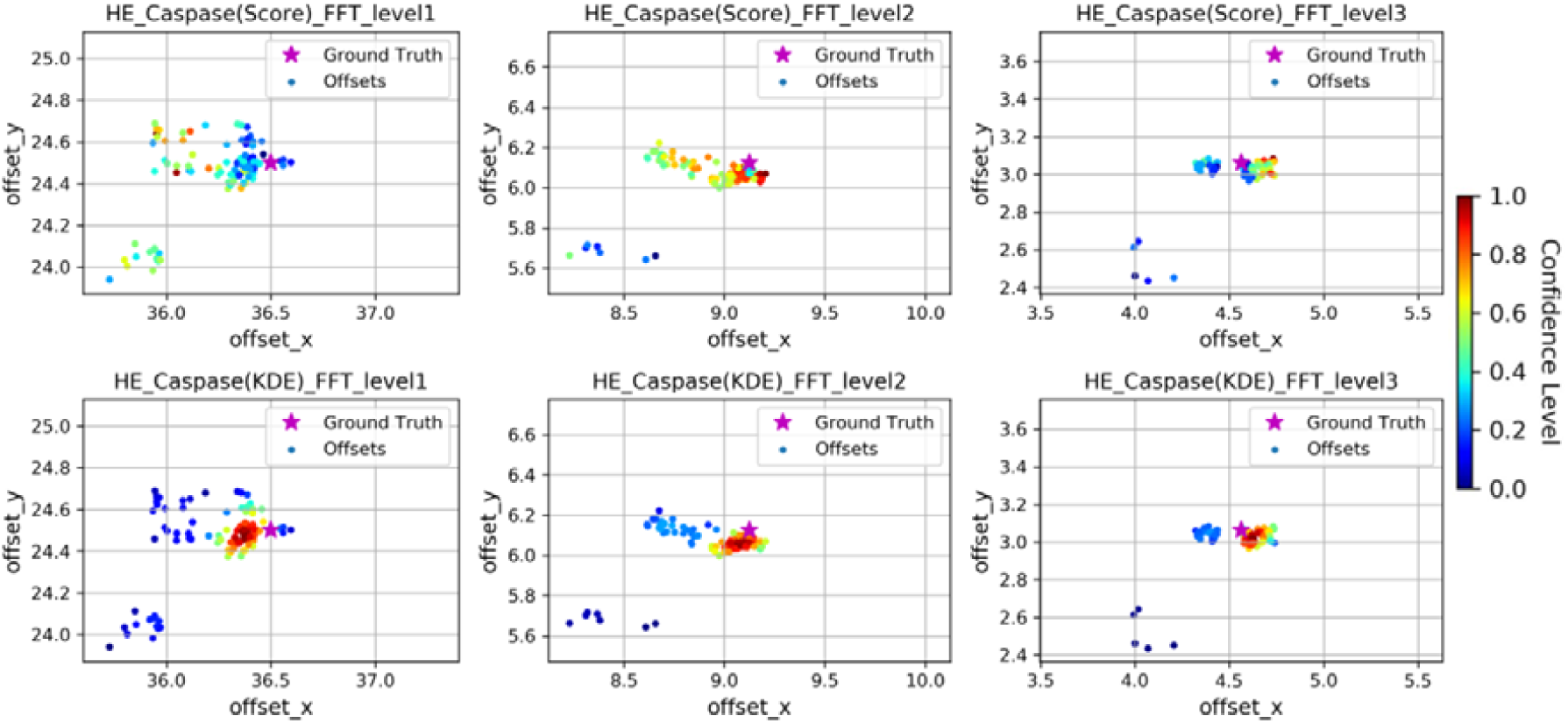
Comparison of KDE and similarity score on reflecting registration success at the patch level. Scatterplots of co-registration offsets at different image levels. Top row: colored with similarity score; Bottom row: colored with KDE. In each subplot, a dot with cold color means registrations with low confidence level, while a dot with hot color means registrations with high confidence level.

### 3. Hierarchical Resolution Regression

As shown in **Figure 2**, the error in offset prediction is strongly correlated with level being viewed. Since image offsets across WSI levels obey a linear downsampling strategy to build their intrinsic pyramidal structure, we introduced linear regression to extrapolate the level 0 offsets. **Figure 5**, we draw the registration results of each image level as scatters, and also draw the line to show their weighted linear regressions. For each WSI pair, the FFT-based regression performed well, with the regression based on either the raw image registration scores was not different from the KDE-weighting scheme. However, the same WSI pair using an alternative raw image registration method (SIFT) was more varied and showed substantial improvement when using the KDE weights rather than the scoring weights. In order to statistically evaluate the performance of our methods, we tested the difference in absolute registration error of each of the four methods (ECC, FFT, SIFT, SIFTenh) using only level 0 image patches or our KDE-weighted linear regression (**Figure 6**). The ECC method improved for the y axis error, but not the x-axis (p=0.24, Student’s T-test). For all other methods, a marked decrease in registration error was observed for both x and y dimensions. The ‘t.test’ function in R [19] was used for calculations.

**Figure 5.**
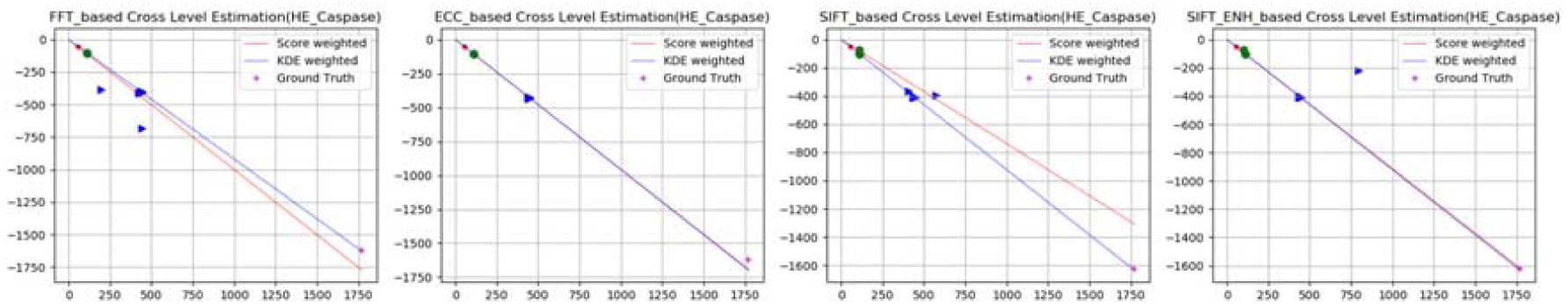
An example of hierarchical linear regression registration. Registration results of each image level were displayed into dots with different shapes for a single WSI. Weighted with similarity score and KDE confidence value, linear regression results are shown in straight lines with different colors. Ground truth is shown in a blue cross.

**Figure 6.**
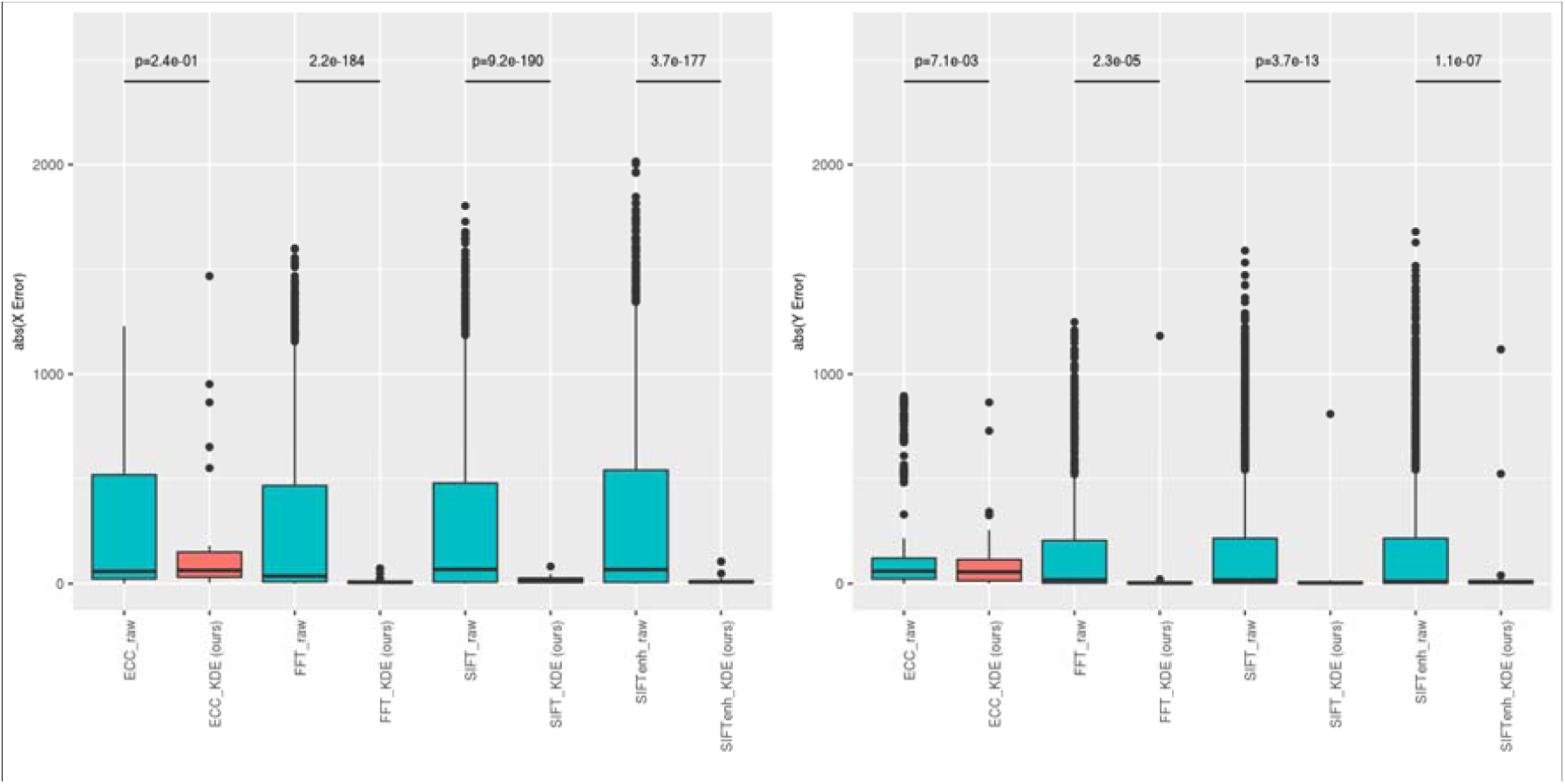
Performance enhancement using the KDE method. All the 30 pairs of WSIs were tested with estimated offsets and ground truth. Registration error was measured with Euclidean distance between the method outputs and ground truth. A Students T-test was used to assess whether the mean absolute erorr in registration was identical between the raw and our KDE-based method. The ideal error profile should be centered as close to zero as possible. Each dot represents the average offset error for each of 30 slides. The left panel is the error in the ‘x’ slide dimension while the right is for the ‘y’ dimension.

## Discussion

In this paper, we proposed an approach that combines traditional rigid image registration methods into a framework that leverages the hierarchical nature of WSIs. We showed the strong advantages of incorporating KDE to better estimate the true offsets compared to the similarity score based approaches. By comparing multiple image registration methods, including FFT, ECC SIFT, and SIFT-ENH, we found that the FFT-based method outperformed than others for our image registration tasks. However, we were able to further increase the precision of the all four registration methods by incorporating our KDE-weighted linear regression

The main limitation of our research is that our method has only been tested on 30 re-stained WSI pairs of H&E and IHC. Consecutive sections are not identical in their cellular structure and composition, thus requiring non-rigid image registration techniques. However, we posit that KDE will still provide a more reliable estimate in this scenario. Furthermore, as our method can be used to match large numbers of image pairs with reliable alignment scores, it provides a scalable framework to collect samples to train a deep neural network, where deep textural patterns beneficial for image alignment can be learned.

The most distinctive part of our work is that we introduced KDE to distill the raw registration result. As most registration methods may return a confidence value for each registration, improper trust in such scores can lead to poor quality results. We show that similarity of the two images after transformation does not always reflect whether the registration was a success or not. Moreover, these approaches do not fully utilize linear relationship across image levels saved in WSI files to ensure maximal accuracy. For example, it is common for many small image patches to be analyzed for a given slide in digital pathology. Each of these patches should have nearly identical x and y offsets. Also, the x and y offsets should follow a fixed linear trend between high and low magnification. By combining the x and y offset estimates across multiple patches from the same slide with the different z magnifications, we have proposed a more robust way to estimate a true x and y offset for each magnification level.

## Authors Contributions

## Acknowledgements

This study is supported by the Division of Laboratory Medicine and Pathology (TJF, SNH) and the Leon Lowenstein Foundation (SNH).

